# Gene Transfer-based Phylogenetics: Analytical Expressions and Additivity via Birth–Death Theory

**DOI:** 10.1101/2022.04.21.489106

**Authors:** Guy Katriel, Udi Mahanaymi, Christoph Koutschan, Doron Zeilberger, Mike Steel, Sagi Snir

**Affiliations:** Department of Mathematics, Ort Braude, Israel; Department of Evolutionary and Environmental Biology, University of Haifa, Israel; RICAM, Austrian Academy of Sciences, Linz, Austria; Department of Mathematics, Rutgers University, USA; School of Mathematics and Statistics, University of Canterbury, NZ

**Keywords:** Genome Dynamics, Markovian Processes, Birth–Death Theory, Phylogenetics, Statistical Consistency

## Abstract

The genomic era has opened up vast opportunities in molecular systematics, one of which is deciphering the evolutionary history in fine detail. Under this mass of data, analysing the point mutations of standard markers is too crude and slow for fine-scale phylogenetics. Nevertheless, genome dynamics events provide far richer information. The *synteny index* (SI) between a pair of genomes combines gene order and gene content information, allowing the comparison of genomes of unequal gene content, together with order considerations of their common genes. Recently, genome dynamics has been modelled as a continuous-time Markov process, and gene distance in the genome as a birth–death–immigration process. Nevertheless, due to complexities arising in this setting such as overlapping neighbourhoods and other confounding factors, no precise and provably consistent estimators could be derived.

Here, we extend this modelling approach by using techniques from birth–death theory to derive explicit expressions of the system’s probabilistic dynamics in the form of rational functions of the model parameters. This, in turn, allows us to infer analytically the expected distances between organisms based on a transformation of their SI. Despite the complexity of the expressions obtained, we establish additivity of this estimated evolutionary distance (a desirable property yielding phylogenetic consistency).

Applying the new measure in simulation studies shows that it attains very accurate results in realistic settings and even under model extensions. In the real-data realm, we applied the new formulation to unique data structure that we constructed - the ordered orthology DB - based on a new version of the EggNOG database, to construct a tree with more than 4.5K taxa. The resulted tree was compared it with a NCBI taxonomy for these organisms. To the best of our knowledge, this is the largest gene-order-based tree constructed and it overcomes flaws found in previous approaches.

## 1 Introduction

The genomic era has reached the point where tasks that seemed imaginary only a decade ago are now within reach. Among these tasks is the inference of the evolutionary history for thousands of species of very close origin. Such a history is depicted in a tree structure and is called a phylogeny. The leaves of that tree correspond to contemporary extant species and the tree’s edges (or branches) represent evolutionary relationships. Despite the impressive advances in the extraction of molecular data, and of ever-increasing quality, finding the underlying phylogenetic tree is still a major challenge that requires reliable approaches for inferring the true evolutionary distances between the species at the tips (leaves) of the tree. The desired tree should preserve the property that the length of the path between any two organisms at its leaves should equal the inferred pairwise distance between these same organisms. When such a tree exists, these distances are called *additive*, as is the distance matrix storing them.

Statistical modelling in which the tree is a parameter of the model are nowadays considered the method of choice for phylogenetic inference. Under this framework, vast efforts have been made, first to model data accurately, and then to draw inferences efficiently from the given data. One such approach is *maximum likelihood* [15,16,17,10,11], where the model (tree) selected is the one maximising the probability of observing the given data.

Standard phylogenetics analyses one or a few ubiquitous genes residing in all species under study, and uses the differences between respective gene copies in order to infer evolution history. Such genes are typically highly conserved by definition and hence cannot provide a strong enough signal to distinguish the shallow branches of the prokaryotic tree. Nevertheless, among prokaryotes, genome dynamics in the form of horizontal gene transfer (HGT) (a mechanism by which organisms transfer genetic material to contemporaneous organisms rather than via vertical inheritance [7,20,24]) and gene loss seem to provide far richer information by affecting both the gene order and gene content. Approaches relying on genome dynamics are mainly divided into gene-order-based and gene-content-based techniques. With the gene-order-based approach [28,14,42], two genomes are considered as permutations of the gene set, and distance is defined as the minimal number of operations needed to transform one genome to the other. The gene-content-based approach [36,40,13] entirely ignores the gene order, and similarity is defined as the size of the set of shared genes. Although a statistical framework has been devised for part of these models [31,41,5,29], to the best of our knowledge, no such framework accounts for HGT.

A related task in this field is the *reconciliation* between a gene tree and the species tree. In this setting, a sequence of events acting on the species tree and yielding the given gene tree is sought. These events may contain events other than HGT while are commonly denoted *duplication, transfer* and *loss* (DTL) [4,37]. These works contain both combinatorial-based approaches such as parsimony [23,8], and model/likelihood-based approaches [38,35]. Neither approach focuses on tree reconstruction, especially reconstructions based on gene order between multiple genes, and therefore, even the underlying evolutionary model they assume is different.

The *synteny index* (SI) [33,1] was suggested as an alternative method to the combinatorial/statistical phylogenetics approaches mentioned above, allowing unequal gene content on one hand while accounting for the order among the shared genes. Here, the locality of a gene in the form of a “neighbourhood” is considered and compared with other genomes. Similarity between genomes is attained by averaging this locality of all the shared genes.

In a recent paper [32], we defined the *jump model* to model genome dynamics, primarily HGT. A genome is defined as a continuous-time Markov process [2]. Under this model, gene distance along the genome can be described as a (critical) birth–death–immigration process. The setting poses intrinsic hurdles such as overlapping neighbourhoods, non-stationarity, confounding factors, and more. Consequently, precise quantities could not be obtained in this earlier work. In particular, basic operations such as gene distance were calculated heuristically.

In this work, we take the jump model and the SI a significant step further by first deriving an analytical expression for the expected time since the divergence between organisms, which is a mandatory step for phylogenetics. Moreover, one of the most important properties in phylogenetics is *model consistency*, which requires that a measure infers accurate distances under a given model of evolution. However, the complexity of the expressions obtained for infering distances do not readily imply consistency of the SI. By using techniques from spectral theory and orthogonal polynomials, we establish in this paper the consistency of the SI measure under the jump model. On the experimental side, we first show that the new mapping provides accurate reconstructions, even for real-life problem sizes and even under an extended jump model that allows gene loss. For real data, we created a new database of ordered orthology groups, based on the EggNOG [18] orthology database, encompassing over 4445 organisms spanning the entire prokaryotic phylogenetic spectrum. Applying the new measure to this database, produces a tree with very high agreement with the NCBI taxonomy [9,30]. To the best of our knowledge, this is the largest genome-dynamics-based tree. In comparison with other SI-based trees, it is evident that the new technique reconstructs significantly more realistic distances, attesting to its capability as a distance measure in other various applications of genome dynamics [6,26].

### Comment

As both the theoretical and the experimental parts are technically involved, we provide in a supplementary text brief self-contained background to the theoretical material employed, as well as further details for the experimental parts.

## 2 Preliminaries

We start by defining a restricted model – *the jump model* – which can be regarded as a transfer between genomes over the same gene set (*equal content*).

### The Jump Model

Let 𝒢^(*n*)^(0) = (*g*_1_, *g*_2_, …, *g*_*n*_) be a sequence of ‘genes’. In our analysis, we will assume that *n* is large enough to allow us to ignore the tips of *𝒢* ^(*n*)^ (or, equivalently, 𝒢^(*n*)^ is cyclic and there are no tips). Consider the following continuous-time Markovian process 𝒢 ^(*n*)^(*t*), *t ≥* 0 on the state space of all *n*! permutations of *g*_1_, *g*_2_, …, *g*_*n*_. Each gene *g*_*i*_ is independently subject to a Poisson process transfer event (at a constant rate *λ*) in which *g*_*i*_ is moved to a different position in the sequence, with each of the possible *n −* 1 positions (between consecutive genes that are different from *g*_*i*_, or at the start or end of the sequence) and with this target location for the transfer selected uniformly at random from these *n −* 1 possibilities.

For example, if *𝒢* ^(*n*)^(*t*) = (*g*_1_, *g*_2_, *g*_3_, *g*_4_, *g*_5_), then *g*_4_ might transfer to be inserted between *g*_1_ and *g*_2_ to give the sequence *𝒢* ^(*n*)^(*t*+*δ*) = (*g*_1_, *g*_4_, *g*_2_, *g*_3_, *g*_5_). The other sequences that could arise by a single transfer of *g*_4_ are (*g*_4_, *g*_1_, *g*_2_, *g*_3_, *g*_5_), (*g*_1_, *g*_2_, *g*_4_, *g*_3_, *g*_5_), and (*g*_1_, *g*_2_, *g*_3_, *g*_5_, *g*_4_). In particular, note that *g*_*i*_ need not necessarily move to a position between two genes; it can also move the the initial or the last position in the sequence.

Since the model assumes a Poisson process, the probability that *g*_*i*_ is transferred to a different position between times *t* and *t* + *δ* is *λδ* + *o*(*δ*), where the *o*(*δ*) term accounts for the possibility of more than one transfer occurring in the time period *δ* (this possibility has probability of order *δ*^2^ and so is asymptotically negligible compared with terms of order *δ* as *δ →* 0). Moreover, a single transfer event always results in a different sequence.

### The Synteny Index

Let *k* be any constant positive integer (note that it may be possible to allow *k* to grow slowly with *n*, but we will not explore such an extension here). For *j ∈ k* + 1, …, *n − k*, the 2*k*-neighbourhood of gene *g*_*j*_ in a genome *𝒢* ^(*n*)^, *N*_2*k*_(*g*_*j*_, *𝒢* ^(*n*)^) is the set of 2*k* genes (different from *g*_*j*_) that have a distance of at most *k* from *g*_*j*_ in *𝒢* ^(*n*)^. We also define *SI*_*j*_(*t*) as the relative intersection size between *N*_*k*_(*g*_*j*_, *𝒢* ^(*n*)^(0)) and *N*_*k*_(*g*_*j*_, *𝒢* ^(*n*)^(*t*)), or formally, 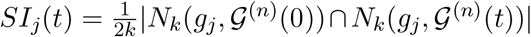 (this is also called *the Jaccard index* between the two neighbourhoods [19]).

Let 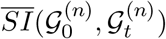 be the average of these *SI*_*j*_(*t*) values over all *j* between *k* + 1 and *n − k*. That is,

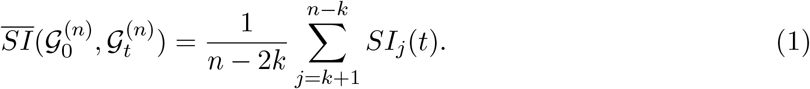

Subsequently, when time *t* does not matter, we simply use 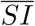 *o*r simply SI where it is clear from the context.

### Phylogenetic Trees and Distances

For a set of species (denoted *taxa*) *𝒳*, a phylogenetic *𝒳* - tree *T* is a tree *T* = (*V, E*) for which there is a one-to-one correspondence between *𝒳* and the set ℒ (*T*) of leaves of *T*. A tree *T* is *weighted* if there is a weight (or length) function associating non-negative weights (lengths) to the edges of *T*. In this paper, we will use the term length, as it corresponds to the number of events or the time span. Edge lengths are naturally extended to *paths*, where the path length is the sum of edge lengths along the path. For a tree *T* over *n* leaves, let *D*(*T*) (or simply *D*) be a symmetric *n×n* matrix where [*D*]_*i,j*_ holds the path length (distance) between leaves *i* and *j* in *T*. A matrix *D*^*′*^ is called *additive* if there is an edge-weighted tree *T* ^*′*^ such that *D*(*T* ^*′*^) = *D*^*′*^. A distance measure *D* is said to be *additive on a model M* if *D* can be transformed (or *corrected*) to give the expected number of events generated under *M*.

### 2.1 Gene Neighbourhood as a Markov Chain

We now introduce a random process, which will play a key role in the analysis of the random variable 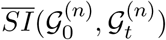. Consider the location of a gene *g*_*i*_, that is not transferred during time period *t*, with respect to another gene *g*_*i*_*′*. Without loss of generality assume *i > i*^*′*^ and let *j* = *i − i*^*′*^. Now, there are *j* ‘slots’ between *g*_*i*_*′* and *g*_*i*_ into which a transferred gene can be inserted, but only *j −* 1 genes in that interval can be transferred. Obviously, a transfer into that interval moves *g*_*i*_*′* one position away from *g*_*i*_, and a transfer from that interval, moves *g*_*i*_*′* closer to *g*_*i*_. This can be modelled as a continuous-time random walk on the state space 1, 2, 3, … with transitions from *j* to *j* + 1 at rate *jλ* (for all *j ≥* 1) and from *j* to *j −* 1 at rate (*j −* 1)*λ* (for all *j ≥* 2), with all other transition rates being 0. This is thus a (generalised linear) birth–death process, illustrated in Fig. 1.

**Fig. 1.**
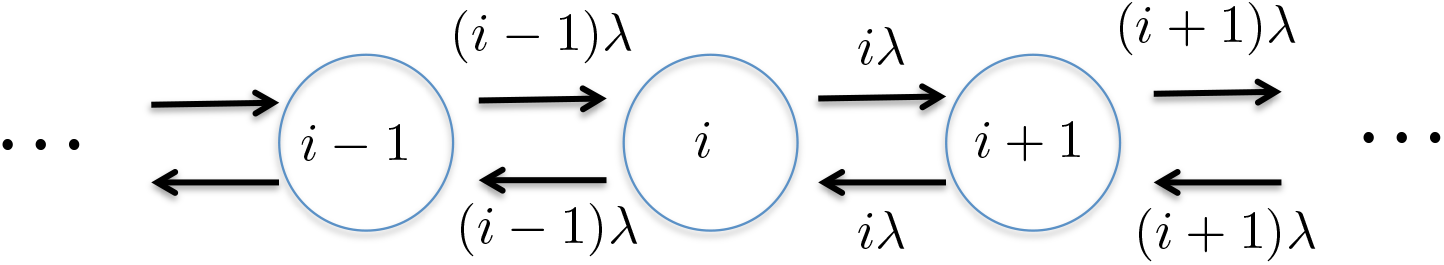
Transitions for the process *X*_*t*_

More formally, we will let *X*_*t*_ denote the random variable that describes the number of slots between two genes under this process described above. Then *X*_*t*_ is a continuous-time random walk on state space 1, 2, 3, …, with an arbitrary initial condition *X*_0_ and transition probabilities of *X*_*t*_ defined as follows:

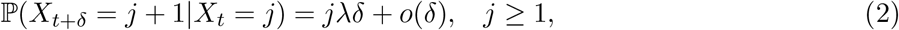

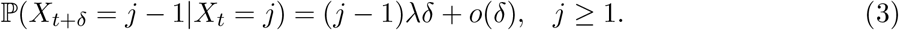

The process *X*_*t*_ is slightly different from the much-studied critical linear birth–death process, for which the rate of birth and death from state *j* are both equal to *j* (here the rate of birth is *j* but the rate of death is *j −* 1), and for which 0 is an absorbing state (here there are no absorbing states). However, this stochastic process is essentially a translation of a critical linear birth–death process with immigration rate equal to the birth–death rate *λ*. This connection is key to the analysis of the divergence times that we establish below.

## 3 Results

### 3.1 Explicit Expressions for the Divergence Time

We now present the main theoretical contribution of this work, which is an analytical expression of divergence times. We first recall a result of [32] that links SI and the transition probabilities of the birth–death process *X*_*t*_. This raises the need to obtain explicit expressions for these probabilities, which we do in Sections 3.2,3.3, making use of known results from the theory of birth–death processes. This theory also allows us (Section 3.4) to give a proof of the monotonicity of the SI as a function of time (in the limit of large *n*), a result that is crucial in order to ensure that we can use our explicit expressions to solve the divergence time in terms of the SI.

Let *p*_*i,j*_(*t*) be the transition probability for *X*_*t*_ to be at state *j*, given that at time 0 it was at state *i*:

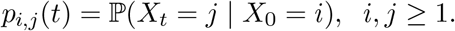

We denote the conditional probability that *X*_*t*_ *∈* [*k*] given that *X*_0_ = *i* by:

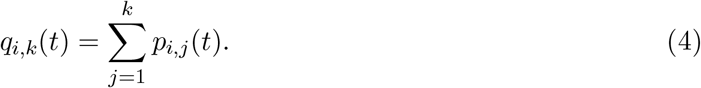

Next, let

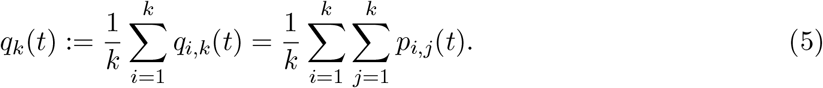

The quantity *q*_*k*_(*t*) is the probability that for a gene at an initial state *i* (i.e., at distance from a reference gene) chosen uniformly at random between 1 and *k*, the process *X*_*\**_ is still between 1 and *k* after time *t*. In [32] we proved the following result:

#### Theorem 1.

*For any given value of t, as n → ∞:*

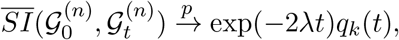

*where* 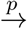 *denotes convergence in probability*.

In the following we assume, without loss of generality, that *λ* = 1 (this is simply rescaling time). The functions *p*_*i,j*_ (*t*) can be expresed as solutions of an infinite system of ordinary differential equations [32] (the Kolmogorov forward equations corresponding to the birth–death process), and these differential equations may be used to numerically approximate *p*_*i,j*_ (*t*) and therefore the key quantity *q*_*k*_(*t*). However, in the present paper we will derive *explicit* algebraic expressions for *p*_*i,j*_ (*t*) and thus *q*_*k*_(*t*). It thereby becomes possible to use Theorem 1 to solve for the divergence time *t* in terms of the SI.

### 3.2 Explicit expressions for *p*_*i,j*_(*t*)

#### Theorem 2.

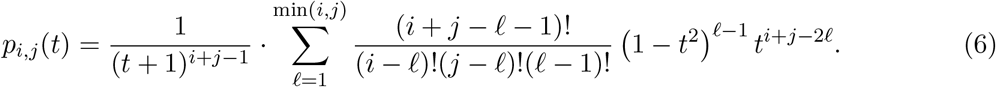

**Proof:** This result follows from some general results for birth–death processes (refer to [3] for these). A simple change in notation will be needed, since the results of [3] involve a birth–death process that is defined on the non-negative integers, whereas our process above i s defined on the positive integers. We therefore define *Y* _*t*_ = *X*_*t*_ *−* 1, so that the process *Y* _*t*_ satisfies an in stance of the general birth–death process described by:

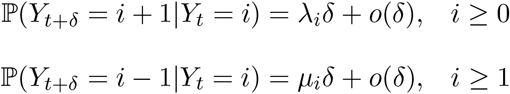

where in our case:

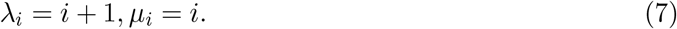

At the heart of the spectral theory of birth–death processes is the Karlin-McGregor representation of the state transition probabilities ([3], Ch. 8, Theorem 2.1):

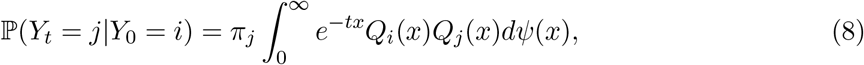

where *dΨ*(*x*) is a measure on [0, *∞*), known as the spectral measure, 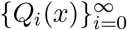 is a sequence of polynomials, orthogonal with respect to the measure *dΨ*, and 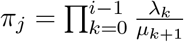.

In the particular case where the birth–death process is given by (7), we have ([3], Ch.8, Eq. 4.14):

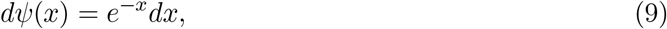

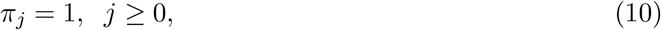

and the polynomials *Q*_*i*_(*x*) are the Laguerre polynomials defined by ([3], Ch.8, Eq. 4.12)

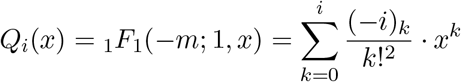

where _1_*F*_1_ is the confluent hypergeometric function, and (*−i*)_*k*_ = (*−i*)(*−i* + 1) · · · (*−i* + *k −* 1). We also have the relation ([3], Ch.8, eq. 4.15)

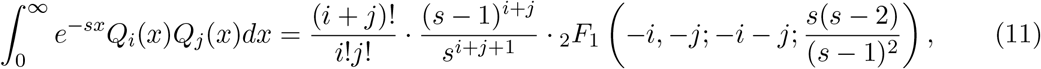

where _2_*F*_1_ is the Gaussian hypergeometric function, defined by:

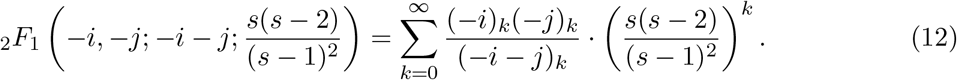

Using (9),(10),(11),(12) with *s* = *t* + 1, (8) leads to ([3], Ch. 8, eq. 4.28):

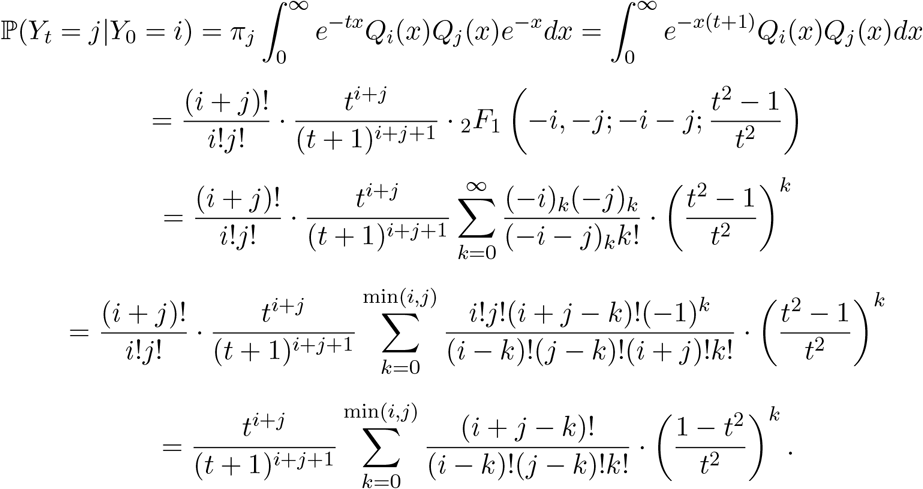

Therefore, going back from the process *Y*_*t*_ to the process *X*_*t*_, we have

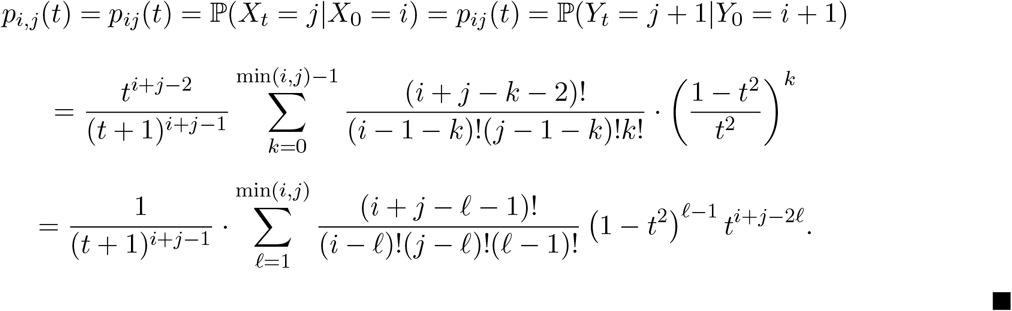

### 3.3 Explicit Expression for *q*_*k*_(*t*)

As stated above, Theorem 1 (originally from [32]) gives an expression for the SI value between two genomes, *𝒢*_0_ and *𝒢*_*t*_. Nevertheless, in that paper, we could not derive an expression only in terms of the number of events that occurred during time *t* (or, alternatively, in a path along the tree of length *λt* “separating” genomes *𝒢*_*i*_ and *𝒢*_*j*_) as we could not derive at an explicit expression for *q*_*k*_. Now that we have obtained explicit expression for *p*_*i,j*_(*t*) in Theorem 2 we can explicitly describe *q*_*k*_ as follows.

#### Theorem 3.

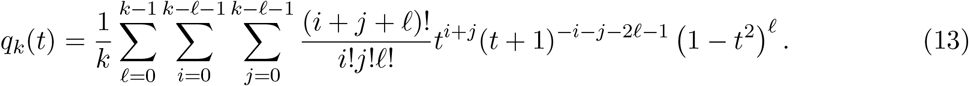

Here are a few instances of the above formula:

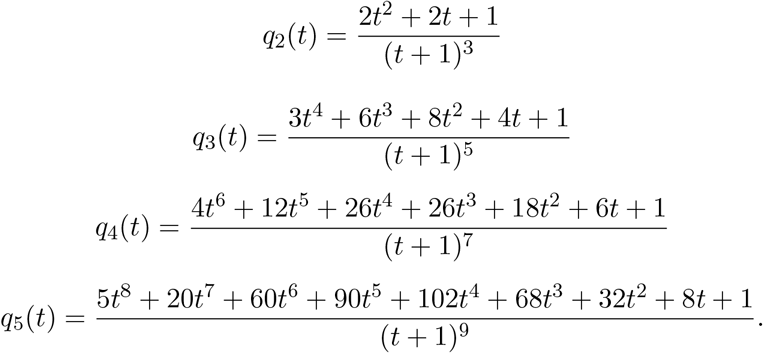

**Proof:** Summing the expressions for *p*_*ij*_(*t*) we get

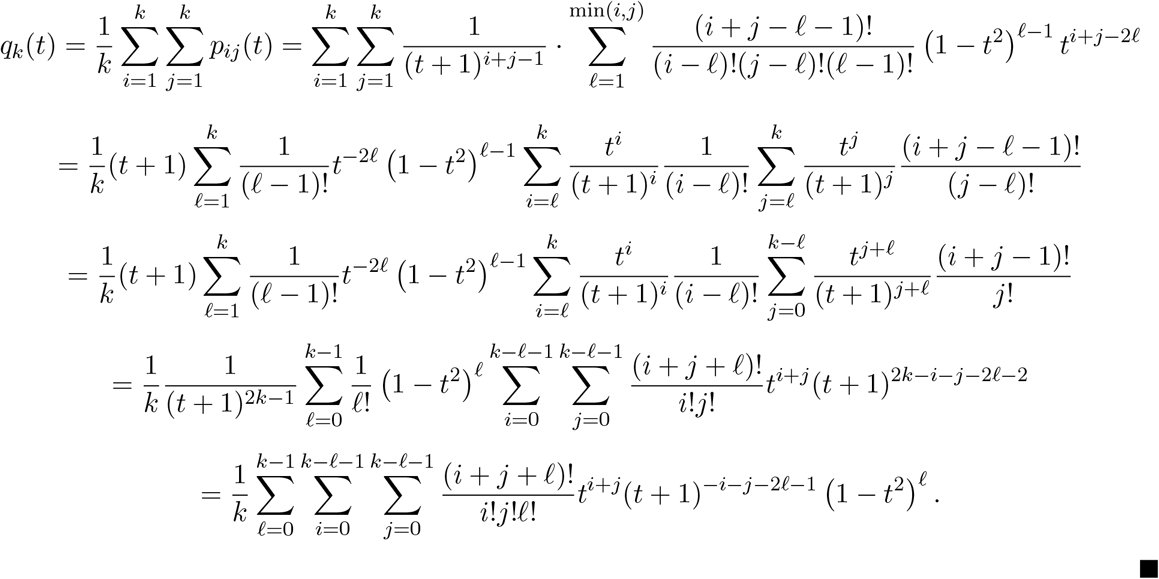

### 3.4 Monotonicity of the SI Measure

Recall that we assumed, without loss of generality, that *λ* = 1, and so our goal now is to prove the monotonicity of the function, *h*_*k*_(*t*) = *e*^*−*2*t*^*q*_*k*_(*t*) and thus (by Theorem 1) the SI measure itself, in the limit of large *n*. In fact we will prove that *q*_*k*_(*t*) itself is monotone decreasing, which obviously implies that *h*_*k*_(*t*) is also monotone decreasing.

#### Theorem 4.

*The function q*_*k*_(*t*) *is monotone decreasing on* [0, *∞*).

Using the representation given by Eq. (8) we have:

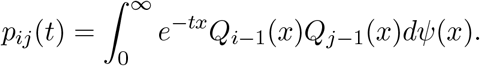

This implies that

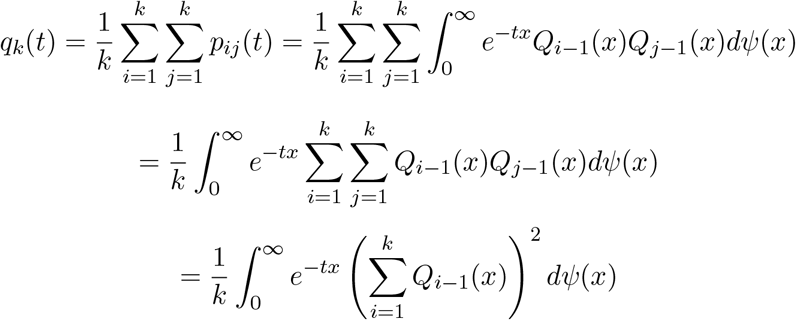

Therefore, by differentiating the above with respect to *t* we obtain:

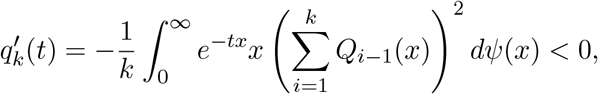

since the integrand is positive. This establishes that *q*_*k*_(*t*) is monotone decreasing.

The fact that *h*_*k*_(*t*) = exp(*−*2*t*)*q*_*k*_(*t*) is strictly monotone decreasing with *t* implies that the inverse function 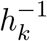 is well-defined. This allows us to use Theorem 1 to reconstruct the time *t* given the SI value and, therefore, estimate the time separating two sequences of genes involving *n* genes (where *n* is large) can be estimated by applying 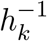 to the 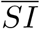 for the two gene sequences. By Theorem 3, we have an explicit value for *h*_*k*_(*t*), so the value 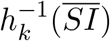 can be calculated by numerically solving a simple equation.

Since the expected number of transfer events is additive on the tree (and proportional to *t*), we can conclude the following:

#### Corollary 1.

*The topology of the underlying unrooted tree T can be reconstructed in a statistically consistent way from the* 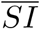 *values by applying the transformation* 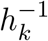, *followed by a consistent distance-based tree reconstruction method such as Neighbour-Joining (NJ)*.

## 4 Experimental Results

In this section, we describe the experiments we conducted to demonstrate the applicability of the theoretical results described above. We begin with simulation results based on the jump model and then move to an analysis of real genomic data.

### 4.1 Simulation results

We simulated the jump model for various values of the number *n* of genes. We set *k* = 3 (i.e. a neighbourhood of 2*k* = 6), the rate was fixed at *λ* = 1 and time *t* varied over the interval [0, 1]. This has yielded a jump probability that was applied to every gene in the initial (*t* = 0) genome. For each value of *t*, the SI between the initial and the final genome was computed. The top part of Fig. 2 displays the value *e*^*−*2*t*^*SI*(*t*) (recall that *λ* = 1 and hence vanishes at the exponent) for each of 10 simulations, and the function *q*_3_(*t*) which is the limit to which *e*^*−*2*t*^*SI*(*t*) converges as *n → ∞*. As can be seen, although there is some variability due to randomness, this variability decreases as *n* increases, and the agreement with the limiting curve *q*_3_(*t*) is clear.

**Fig. 2.**
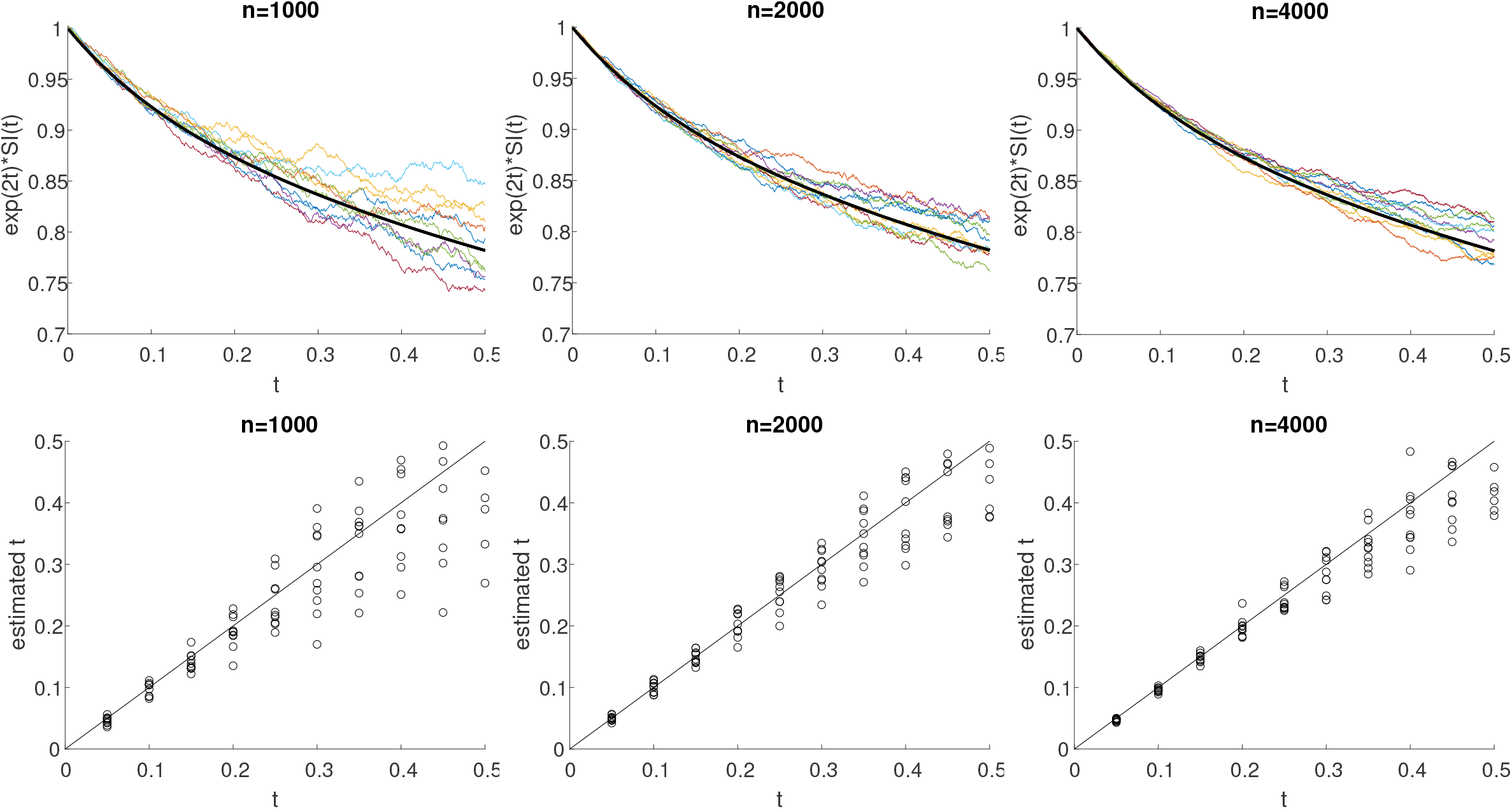
Simulation Results, the pure jump model:. Genome sizes *n* = 1000, 2000, 4000. Top: Comparison of the curves *e*^*λt*^*SI*(*t*) computed using simulation with the limiting curve *q*3(*t*), Bottom: Estimated vs effective *t*.

In a related experiment, we checked how well the value *SI*(*t*), computed using the simulated data, can be used to estimate the time *t*. For each value of *t*, we compute *SI*(*t*) from the simulated data, and use this to estimate *t* by numerically solving the following equation:

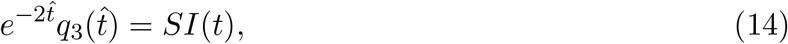

In the lower part of Fig. 2 the true value of *t* is compared with the estimated values 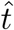 for 10 simulations.

We note that the relevant values of *λt* as found in [32] are around 0.4 for distances within the phylogenetic rank of genus. We see that the error is almost insignificant even for realistic genome sizes, as we have here.

Next, we extended the pure jump model to include gain/loss events, both occurring with a probability *p*, so the expected genome length is fixed. We again used Eq. (13) to infer the distances. The results for the gain/loss probabilities 0.1 and 0.2 are shown in Fig. 3. As can be seen, although the estimation is less accurate than in the pure jump model, the lag in estimated 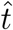 is linear with *t* and maintains additivity (as noted in [32] under the gain/loss model).

**Fig. 3.**
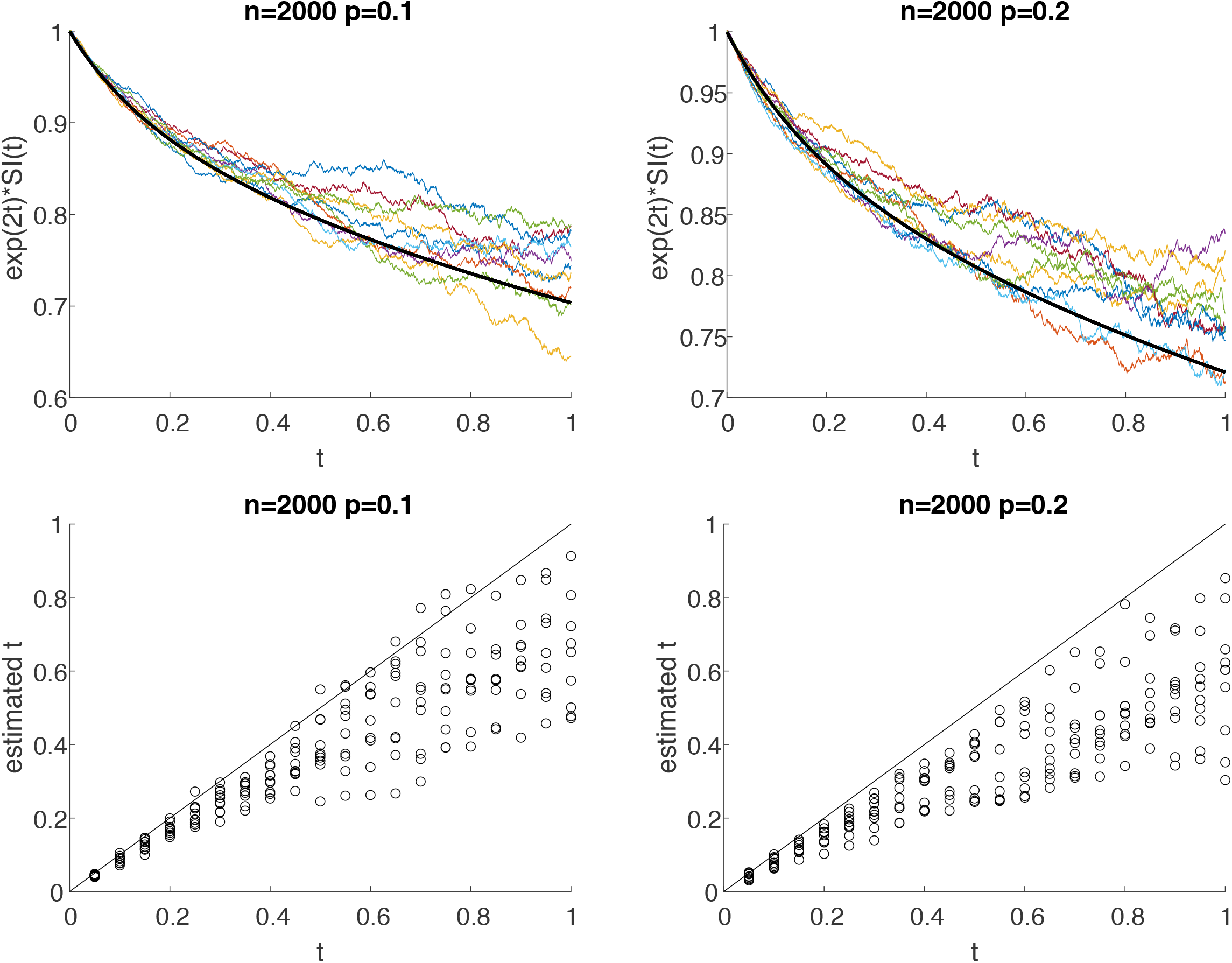
Simulation Results, Jump plus gain/loss:. genome size *n* = 2000 Top: Estimated vs effective 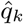 for gain/loss *p* = 0.1, 0.2. Bottom: Estimated vs effective 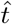 or gain/loss *p* = 0.1, 0.2.

### 4.2 Real Data Results

Here we report the real data results obtained using the new technique. Because of space limitations, and for the sake of reconstructability, fuller details and data are provided in the supplementary text and material respectively. We applied our method to real genomic data consisting of 4445 prokaryotes taken from the orthology data base EggNOG [18] with 4.4M clusters of orthologous groups (COGs) [39]. For each COG, EggNOG provides a flat ‘members’ file indicating the organisms that harbour this gene, along with its location in the genome. This allowed us to sort the genes by location along the genome. Within this representation, a genome is simply a list of COGs sorted by genome location, where the COG names are universal across all organisms. Hence, we can infer neighbourhood similarities across genomes and therefore the pairwise SI values between any two genomes which we then store in an *n × n* SI matrix. We set *k* = 10 which was found to be informative for these data [33,32] and computed SI for all pairs of taxa. The crude SI values are strongly concentrated around 0.02, as shown in Fig. 4(R). In order to convert the SI values to a dissimilarity measure, we set *d*_*SI*_ = 1 *− SI*. Once a (pairwise) dissimilarity *D* matrix has been computed, we can then apply a distance-based phylogenetic method to estimate a tree *T* in which the leaves are labelled by the organisms under study.

**Fig. 4.**
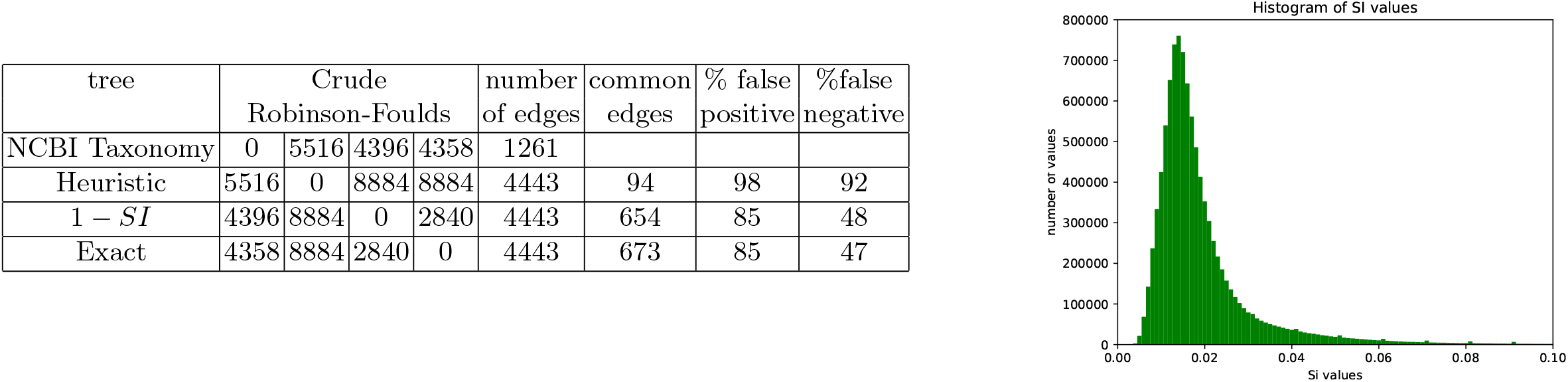
**Left:** Robinson–Foulds distances. **Right:** SI values.

Path distances between the leaves of *T*, should approximate the distances in *D*. The most accurate algorithm for this task is the neighbour joining (NJ) algorithm [27]. Therefore, we used the program Neighbour from Phylip [12] to construct a tree that we call the 1 *− SI* tree. Recall now that Eq. (14) was devised to “correct” the crude *d*_*SI*_ and provide a (provably) more reliable distance. Hence, we “corrected” the SI matrix accounting to Eq. (14) (specifically, finding 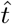 by solving Eq. (14) for the appropriate SI value in the matrix) and then applied Neighbour to this matrix, yielding the *exact tree*. Finally, as in [32], we did not have an explicit expression for distance and were forced to develop a simulation-based heuristic, we also constructed the *heuristic tree* by using Formula (9) from [32].

EggNOG labels its organisms with the same taxon ID used by the NCBI taxonomy database [9]. This database is also furnished with taxonomic ranks in a child-to-parent relationship that we can use for our task. We therefore constructed a tree from this child–parent relationship. This NCBI tree spans about 1.2M organisms with maximum depth (i.e. ranks) of 39. We extracted the tree induced by EggNOG 4445 taxa^1^ and used this tree as a reference tree, dubbed *the NCBI tree*. The four trees appear in Fig. 5 in two formats - rectangular (L) and polar(R). As can be seen, the 1 *− SI* and the heuristic trees exhibit serious flaws we will elaborate on later.We wanted to measure the distance from each of the three reconstructed trees to the reference NCBI tree. A common tree distance measure is the Robinson–Foulds (RF) symmetric difference [25], which can be used to derive the false positive and false negative (FP, FN) rates. The relevant distances are presented in Table 4(L). As can be seen, the exact tree from Eq. (14) is the most similar to the NCBI tree and the heuristic tree is the least similar.

**Fig. 5.**
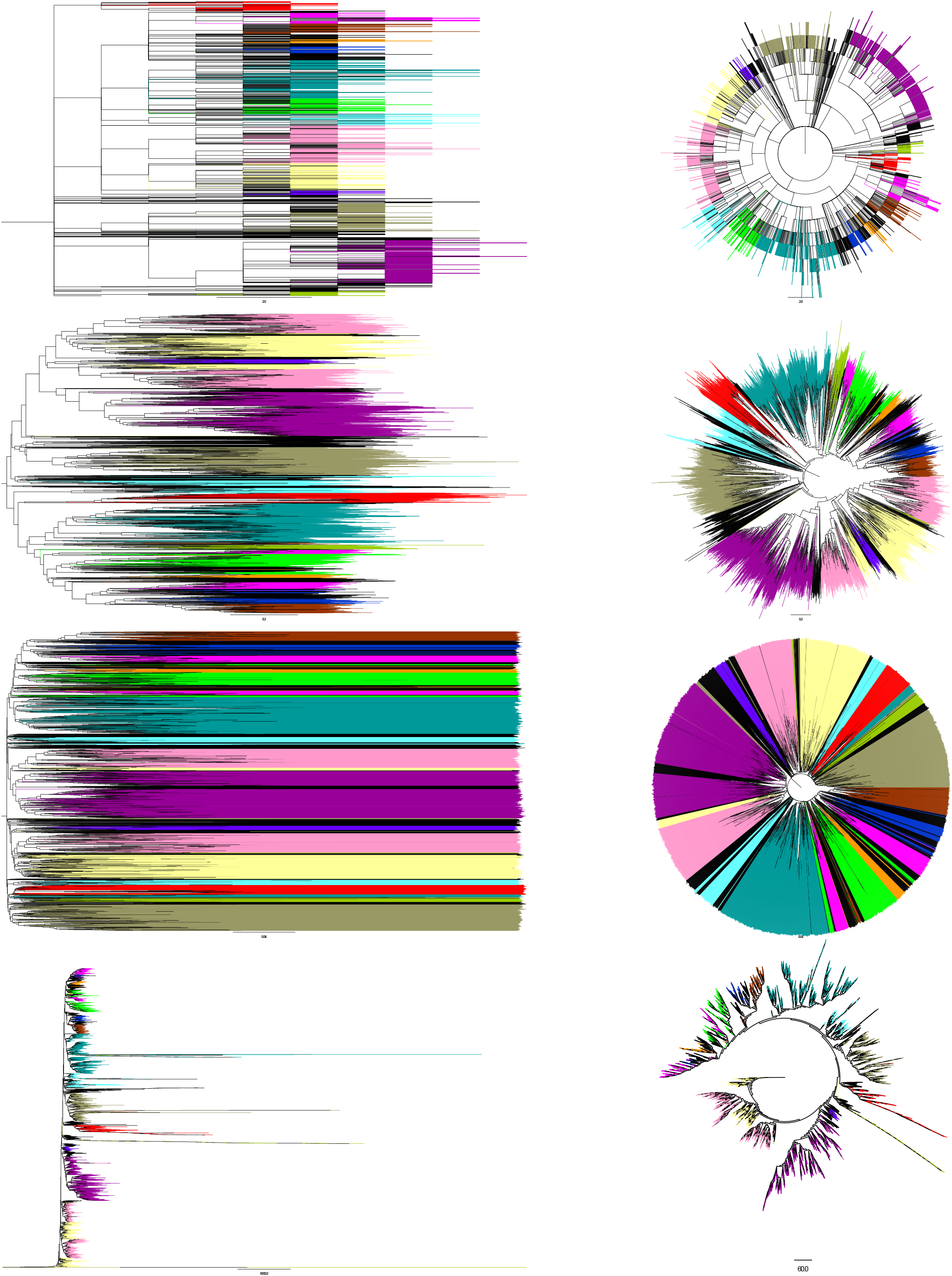
Coloured Trees: Left: rectangular shape; right: polar shape. From the top: (1) The NCBI Taxonomy, (2) The exact SI tree, (3) the 1 *− SI* tree, (4) the heuristic exp. decay tree (the polar shape on right has log distances to accommodate the extremely long branches).

The RF distance is very sensitive and uninformative for large trees [34]. Hence, we adopted a compatibility measure to allow a more intuitive assessment. We divided the tree into disjoint subtrees with sizes of between 80 and 800 taxa, resulting in 14 subtrees in total. This tree partitioning served as a *reference colouring* where each such subtree was mapped to a colour and all taxa (leaves) in a subtree had the same colour. For a colour *c*, the *c*-subtree is defined as the minimal connected graph containing all *c*-coloured nodes. Given a coloured tree (i.e. with some of the nodes coloured), such a colouring is said to be *convex* on that tree, if for every two colours *c* and *c*^*′*^, the *c*- and *c*^*′*^-coloured subtrees are disjoint [22,21]. It is clear that the NCBI tree is convex, since the colouring is defined by this tree, i.e. for disjoint subtrees. Nevertheless, we aimed to test how far from convexity the NCBI colouring on the other trees is. There are rigorous definitions for the latter (the *recoloring distance [22]*); however, we used this approach to provide an intuitive and visual measure of compatibility, as demonstrated in Fig. 5.

As can be seen from the figure, all three trees maintained decent convexity under the NCBI colouring; however, it seems the exact tree has fewer violations than the heuristic and the 1 *− SI* trees. Fig. 5 also reveals major flaws in the heuristic and the 1 *− SI* approaches that are corrected by the exact approach. The 1 *− SI* approach takes crude values as the distances. These values are excessively concentrated around a tiny value of 0.02, causing severely distorted branch lengths, resulting in an artificially ultrametric tree with extremely short internal branches (third row in Fig. 5), which may disappear under bootstrapping, yielding a poorly resolved tree. Alternatively, the heuristic approach of [32], apart from achieving an outstandingly high RF distance, produces few exceptionally long branches non-proportional to the rest of the branches (left tree in the fourth row in Fig. 5). To conclude this part, our real-data experiments showed that the theoretical conversion achieves its goal by producing a realistic distance, thereby correcting the severe flaws caused by the two other approaches.

## 5 Conclusions

In this paper, we explored the consequences of modeling genome organisation as a continuous-time Markov process. Although the initial modelling was suggested recently, fundamental problems were left open, making it impossible to formally answer basic questions such as the time since divergence on a tree or the additivity of the synteny index as a phylogenetic marker. Here, we have significantly advanced this front by applying advanced mathematical tools from analysis and algebra to arrive at a rational function describing the transition probabilities, and the use of spectral theory and orthogonal polynomials, to prove the measure’s consistency.

In the experimental realm, we presented accurate results for the new analytic expressions even in real-life genome sizes and event rates. For the real data analysis, we built an ordered database of orthologous groups across 4445 prokaryotes, to which we applied our measure. To the best of our knowledge, there is no such database of this size in terms of orthologous groups or the number of taxa. Such a database could have multiple uses, apart from phylogenetics.

Applying our new measure to this database produced a tree that was in high accordance with the NCBI taxonomy for these organisms. Importantly, the new measure reconstructed realistic distances, as opposed to the previous measures, even the heuristic measure that was developed based on simulations. Reconstructing accurate distances has prime importance for establishing the jump model as an underlying model of genome dynamics.

We expect that both the rigour developed here for the modelling and the data resources will be instrumental in further analyses of other genome architectures such as operon and pseudogene formation.

## Footnotes

1 This was done by removing the leaves that were not in the selection, and the paths leading solely to them.

## References

1. O. Adato, N. Ninyo, U. Gophna, and S. Snir. Detecting horizontal gene transfer between closely related taxa. PLoS computational biology, 11(10):e1004408, 2015.

2. L. J. Allen. An introduction to stochastic processes with applications to biology. Chapman and Hall/CRC, 2010.

3. W. J. Anderson. Continuous-time Markov chains: An applications-oriented approach. Springer Science & Business Media, 2012.

4. M. S. Bansal, M. Kellis, M. Kordi, and S. Kundu. Ranger-dtl 2.0: rigorous reconstruction of gene-family evolution by duplication, transfer and loss. Bioinformatics, 34(18):3214–3216, 2018.

5. P. Biller, L. Guéguen, and E. Tannier. Moments of genome evolution by double cut-and-join. BMC bioinformatics, 16(14):S7, 2015.

6. D. Che, G. Li, F. Mao, H. Wu, and Y. Xu. Detecting uber-operons in prokaryotic genomes. Nucleic acids research, 34(8):2418–2427, 2006.

7. W. F. Doolittle. Phylogenetic classification and the universal tree. Science, 284(5423):2124–2128, 1999.

8. J.-P. Doyon, C. Scornavacca, K. Y. Gorbunov, G. J. Szöllősi, V. Ranwez, and V. Berry. An efficient algorithm for gene/species trees parsimonious reconciliation with losses, duplications and transfers. In RECOMB International Workshop on Comparative Genomics, pages 93–108. Springer, 2010.

9. S. Federhen. The NCBI Taxonomy database. Nucleic Acids Research, 40(D1):D136–D143, 12 2011.

10. J. Felsenstein. Cases in which parsimony or compatibility methods will be positively misleading. Systematic zoology, 27(4):401–410, 1978.

11. J. Felsenstein. Evolutionary trees from dna sequences: a maximum likelihood approach. Journal of molecular evolution, 17(6):368–376, 1981.

12. J. Felsenstein. Phylip (phylogeny inference package), version 3.5 c. 1993.

13. S. T. Fitz Gibbon and C. H. House. Whole genome-based phylogenetic analysis of free-living microorganisms. Nucleic acids research, 27(21):4218–4222, 1999.

14. S. Hannenhalli and P. A. Pevzner. Transforming cabbage into turnip: polynomial algorithm for sorting signed permutations by reversals. volume 46, pages 1–27. ACM, 1999.

15. M. D. Hendy and D. Penny. A framework for the quantitative study of evolutionary trees. Systematic zoology, 38(4):297–309, 1989.

16. M. D. Hendy and D. Penny. Spectral analysis of phylogenetic data. Journal of classification, 10(1):5–24, 1993.

17. M. D. Hendy, D. Penny, and M. Steel. A discrete fourier analysis for evolutionary trees. Proceedings of the National Academy of Sciences, 91(8):3339–3343, 1994.

18. J. Huerta-Cepas, D. Szklarczyk, D. Heller, A. Hernández-Plaza, S. K. Forslund, H. Cook, D. R. Mende, I. Letunic, T. Rattei, L. Jensen, C. von Mering, and P. Bork. eggNOG 5.0: a hierarchical, functionally and phylogenetically annotated orthology resource based on 5090 organisms and 2502 viruses. Nucleic Acids Research, 47(D1):D309–D314, 11 2018.

19. P. Jaccard. Étude comparative de la distribution florale dans une portion des alpes et des jura. Bull Soc Vaudoise Sci Nat, 37:547–579, 1901.

20. E. V. Koonin, K. S. Makarova, and L. Aravind. Horizontal gene transfer in prokaryotes: quantification and classification. Annual Reviews in Microbiology, 55(1):709–742, 2001.

21. S. Moran and S. Snir. Efficient approximation of convex recolorings. Journal of Computer and System Sciences (JCSS), 73:1078–1089, 2007. Earlier version appeared in APPROX/RANDOM 2005.

22. S. Moran and S. Snir. Convex recolorings of strings and trees: Definitions, hardness results and algorithms. J. Comput. Syst. Sci., 74(5):850–869, 2008.

23. L. Nakhleh, D. Ruths, and L.-S. Wang. Riata-hgt: a fast and accurate heuristic for reconstructing horizontal gene transfer. In International Computing and Combinatorics Conference, pages 84–93. 2005.

24. H. Ochman, J. G. Lawrence, and E. A. Groisman. Lateral gene transfer and the nature of bacterial innovation. nature, 405(6784):299, 2000.

25. D. F. Robinson and L. R. Foulds. Comparison of phylogenetic trees. Mathematical biosciences, 53(1-2):131–147, 1981.

26. I. Rogozin, K. Makarova, J. Murvai, E. Czabarka, Y. Wolf, R. Tatusov, L. Szekely, and E. Koonin. Connected gene neighborhoods in prokaryotic genomes. Nucleic Acids Res, 30:2212–2223, 2002.

27. N. Saitou and M. Nei. The neighbor-joining method: a new method for reconstructing phylogenetic trees. Molecular Biology and Evolution, 4(4):406–425, 1987.

28. D. Sankoff. Edit distance for genome comparison based on non-local operations. In Annual Symposium on Combinatorial Pattern Matching, pages 121–135. Springer, 1992.

29. D. Sankoff and J. H. Nadeau. Conserved synteny as a measure of genomic distance. Discrete applied mathematics, 71(1-3):0247–257, 1996.

30. C. L. Schoch, S. Ciufo, M. Domrachev, C. L. Hotton, S. Kannan, R. Khovanskaya, D. Leipe, R. Mcveigh, K. O’Neill, B. Robbertse, S. Sharma, V. Soussov, J. P. Sullivan, L. Sun, S. Turner, and I. Karsch-Mizrachi. NCBI Taxonomy: a comprehensive update on curation, resources and tools. Database, 2020, 08 2020. baaa062.

31. S. Serdoz, A. Egri-Nagy, J. Sumner, B. R. Holland, P. D. Jarvis, M. M. Tanaka, and A. R. Francis. Maximum likelihood estimates of pairwise rearrangement distances. Journal of theoretical biology, 423:31–40, 2017.

32. G. Sevillya, D. Doerr, Y. Lerner, J. Stoye, M. Steel, and S. Snir. Horizontal Gene Transfer Phylogenetics: A Random Walk Approach. Molecular Biology and Evolution, 37(5):1470–1479, 12 2019.

33. A. Shifman, N. Ninyo, U. Gophna, and S. Snir. Phylo si: a new genome-wide approach for prokaryotic phylogeny. Nucleic acids research, 42(4):2391–2404, 2013.

34. K. Siu-Ting, D. Pisani, C. J. Creevey, and M. Wilkinson. Concatabominations: Identifying Unstable Taxa in Morphological Phylogenetics using a Heuristic Extension to Safe Taxonomic Reduction. Systematic Biology, 64(1):137–143, 09 2014.

35. J. Sjöstrand, A. Tofigh, V. Daubin, L. Arvestad, B. Sennblad, and J. Lagergren. A bayesian method for analyzing lateral gene transfer. Systematic biology, 63(3):409–420, 2014.

36. B. Snel, P. Bork, and M. A. Huynen. Genome phylogeny based on gene content. Nature genetics, 21(1):108, 1999.

37. M. Stolzer, H. Lai, M. Xu, D. Sathaye, B. Vernot, and D. Durand. Inferring duplications, losses, transfers and incomplete lineage sorting with nonbinary species trees. Bioinformatics, 28(18):i409–i415, 2012.

38. G. J. Szöllősi, E. Tannier, N. Lartillot, and V. Daubin. Lateral gene transfer from the dead. Systematic biology, 62(3):386–397, 2013.

39. R. L. Tatusov, D. A. Natale, I. V. Garkavtsev, T. A. Tatusova, U. T. Shankavaram, B. S. Rao, B. Kiryutin, M. Y. Galperin, N. D. Fedorova, and E. V. Koonin. The cog database: new developments in phylogenetic classification of proteins from complete genomes. Nucleic acids research, 29(1):22–28, 2001.

40. F. Tekaia and B. Dujon. Pervasiveness of gene conservation and persistence of duplicates in cellular genomes. Journal of molecular evolution, 49(5):591–600, 1999.

41. L.-S. Wang and T. Warnow. Estimating true evolutionary distances between genomes. In Proceedings of the thirty-third annual ACM symposium on Theory of computing, pages 637–646. ACM, 2001.

42. S. Yancopoulos, O. Attie, and R. Friedberg. Efficient sorting of genomic permutations by translocation, inversion and block interchange. Bioinformatics, 21(16):3340–3346, 2005.

