## Supplementary Text for "Gene Transfer-based Phylogenetics: Analytical Expressions and Additivity via Birth–Death Theory"

### Supplementary text for manuscript "Gene Transfer based Phylogenetics: Analytical Expressions and Additivity via Birth-Death Theory"

April 18, 2022

#### 1 Brief relevant theoretical background

In this section we provide brief theoretical background regarding stochastic processes that is required to the undestanding of material in the main paper body. This material is mostly taken from the text books of Feller [3], Grimmett and Stirzaker [4] and Allen [1]. We note that the following is not intended to replace the formal comprehensive text in a standard text book.

##### 1.1 Poisson processes

A *Poisson process* with *rate* (or *intensity*) $\lambda$ is a process generating events in time, such that for sufficiently small $h$, the probability for an event between $t$ and $t+h$ is $h\lambda + o(h)$ and for more than a single event $o(h)$. It can be shown that the probability distribution (for number of events $k$ during time period $t$) of a Poisson process with rate $\lambda$ is:

$$\mathbb{P}(k, \lambda t) = \frac{(\lambda t)^k}{k!} e^{-\lambda t}. \tag{1}$$

A special case of this distribution that we use in the main text is the probability of "no event":

$$\mathbb{P}(0, \lambda t) = e^{-\lambda t}. \tag{2}$$

##### 1.2 Simple birth-death processes

A *simple birth process* with rate $\lambda$ is a Poisson process with $n$ individuals, each giving birth to a new individual with probability $\lambda h + o(h)$ in the time interval $(t, t+h)$. Under a *simple death proces*, each individual dies at rate $\mu$.

#### 1.3 Continuous-time Markov chains

**Definition 1** *[[1]5.1] A stochastic process* $\{X(t) : t \in [0, \infty)\}$ *such that* $X(t) \in \{1, 2, 3, \ldots\}$*, is called a* continuous-time Markov chain *(CTMC) if it satisfies the following condition: For any sequence of real numbers satisfying* $0 \le t_0 < t_1 < \ldots < t_n < t_n + 1$*, the probability to be in time* $t_{n+1}$ *at any state* $i_{n+1}$ *given all the previous states, dependes only at the state at time* $t_n$*.*

The dependence only on the current step is denoted the *Markov property* and is extremely common and useful in stochastic processes. It can be readily seen that a birth-death process is a continuous-time Markov chain where the size of the population $n$ is state set $X(t)$, and any new event moves to new state. Due to the Markov property, it is a convention to denote by $p_{i,j}(t)$ the probability to move from a state $i$ to $j$ within a time interval $t$. Therefore, in a birth-death process, the birth-death rate at state $n$ is $n(\lambda - \mu)$. At state $n = 0$, a rate zero is obtained and then 0 is an *absorbing state*. Such a process is called *linear* as the increment in each state change is constant, and is *critical* when $\lambda = \mu$, and *supercritical* and *subcritical* for $\lambda > \mu$ and $\lambda < \mu$ respectively. An *immigration process* is a Poisson process with a constant rate $\nu$ independent of the population size $n$. It is easy to see that under immigration, no absorbing state exists and the compound rate is $n(\lambda - \mu) + \nu$.
The *forward Kolmogorov differential equations* expresses the change in $p_{i,j}(t)$ with time:

$$\frac{dp_{ij}(t)}{dt} = \lambda_{j-1} p_{i,(j-1)}(t) - (\lambda_j + \mu_j) p_{ij}(t) + \mu_{j+1} p_{i(j+1)}(t). \tag{3}$$

Under the linear birth-death process, we have $\lambda_j = j\lambda$ and $\mu_j = j\mu$ for every $j$ yielding:

$$\frac{dp_{ij}(t)}{dt} = (j-1)\lambda p_{i,(j-1)}(t) - j(\lambda + \mu) p_{ij}(t) + (j+1)\mu p_{i(j+1)}(t). \tag{4}$$

#### 1.4 The spectral theory of Birth-death processes

Birth-death processes are a special type of continuous-time Markov chain, in which the stste space is the set of non-negative integers, and the only allowed transitions are among adjacent integers. The state $Y_t$ of such a process therefore satisfies

$$P(Y_{t+\Delta t} = i + 1 | Y_t = i) = \lambda_i \Delta t + o(\Delta t), \quad i \ge 0$$

$$P(Y_{t+\Delta t} = i - 1 | Y_t = i) = \mu_i \Delta t + o(\Delta t), \quad i \ge 0,$$

where $\lambda_i, \mu_i$ are called the birth and death rates, respectively. The special structure of birth-death processes enables a powerful theory, known as the spectral theory of birth-death processes [5]. For a detailed exposition of this theory we refer to [2], Ch. 8. This theory provides a representation of the state probabilities of the process:

$$P(Y_t = j|Y_0 = i) = \pi_j \int_0^\infty e^{-tx} Q_i(x) Q_j(x) d\psi(x), \;\; i, j \geq 0 \tag{5}$$

known as the Karlin-McGreggor representation. Here $\{Q_i(x)\}_{i=0}^\infty$ is a sequence of polynomials ($Q_i(x)$ has degree $i$), defined by the recursion:

$$Q_0(x) = 1$$

$$-xQ_0(x) = -(\lambda_0 + \mu_0)Q_0(x) + \lambda_0 Q_1(x)$$

$$-xQ_i(x) = \mu_i Q_{i-1}(x) - (\lambda_i + \mu_i)Q_i(x) + \lambda_i Q_{i+1}(x), \quad i \geq 1,$$

and the integral in (5) is a Stieltjes integral with respect to a probability distribution function $\psi(x)$ associated to the polynomial sequence, known as the spectral measure, with respect to which the polynomials are orthogonal, that is:

$$\pi_j \int_0^\infty e^{-tx} Q_i(x) Q_j(x) d\psi(x) = \delta_{ij},$$

($\delta_{ij} = 1$ if $i = j$ and $\delta_{ij} = 0$ otherwise), and the coefficients $\pi_j$ are given by $\pi_0 = 0$ and

$$\pi_j = \frac{\lambda_0 \cdot \lambda_1 \cdots \lambda_{j-1}}{\mu_1 \cdot \mu_2 \cdots \mu_j}, \quad j \geq 1.$$

While the representation (5) is valid for an arbitrary birth-death processes, only special cases in which the dependence of the coefficients $\lambda_i, \mu_i$ on $i$ is particularly simple allow finding explicit expressions for the polynomials $Q_i(x)$, the spectral measure $\psi$, and hence the transition probabilities given by (5). Specifically, this is the case for the coefficients which are of interest in this work, given by:

$$\lambda_i = i + 1, \mu_i = i. \tag{6}$$

In this case the spectral measure is an exponential distribution, the orthogonal polynomials are Laguerre polynomials, and the formula (5) provides exlicit expressions for the transitions probabilities, as given in the main text. The Karlin-McGreggor representation (5) also provides the key tool for proving the monotonicity of the SI measure $q_k(t)$.

### 2 Detailed Experimental Study

#### 2.1 The NCBI Taxonomy Hierarchy

The NCBI Taxonomy database [10] contains 1,241,055 species (2020), in a hierarchical structure. The tree structure in a Newick (text) format can be found at suppMaterial/NCBIFullTree.nwk file. This tree has 1,119,775 leaves, orgenized in 39 level where the root level is considered as level one. The tree is highly unbalanced and unresolved: The root of this tree has 5 children with one of these children has 1,103,368 offspring - almost the entire taxa set. The maximal number of children for a node is 41516. Figure 1(.1) depicts the number of leaves and nodes in any level.

#### 2.2 Constructing the Ordered Orthology DB from the EggNOG Repository

TheEggNOG 5.0 D.B. database contains information over 4,445 prokaryotes with average of 3624.3 genes per genome, and a minimum and a maximum of 359 and 11,511 genes per genome respectively. In order to build our ordered orthology DB as described in the main text, we relied on the following data sources of EggNOG (see a schematic structure at Figure 1(.2)). A set of GBFF files, one or more for each genome indicate the location and the nucleotide sequence of each gene in the genome. A map file maps genome names to their respective GBFF file. Next there is a single "members" file that associate each gene in each genome, with its corresponding COG (Cluster of Orthologous Groups) name. This system of files and pointers, enables to construct the EggNOG-based Ordered Orthology DB, in which every genome is represented as an ordered list of its constituting gene, and each gene is represented by its COG name.

#### 2.3 The NCBI Derived subTree

In order to restrict the NCBI taxonomy hierarchy to our 4,445 genomes from EggNOG, we pursued the following the following steps. These genomes constitute the leaves of this subtree. We created this subtree from the original NCBI tree by a pruning process that is described later. The mean length of a genome of these 4,445 bacteria is 3624.3 genes, with minimal genome length of 359 genes, and maximal genome Length of 11,511 genes. The induced bacteria taxonomy subtree holds 5,705 nodes in 11 levels; Its root is node 237368 - 'Candidatus Scalindua brodae', from level 3 of the original NCBI tree. The number of leaves and nodes in any level, appears in the Figure 1(.3). The tree structure as text format is at suppMaterial/EggNOGFullTree.txt file, and in newick format is at suppMaterial/EggNOGFullTree.nwk file.

| Level | #leaves | #nodes |
|---|---|---|
| 1 | 0 | 1 |
| 2 | 0 | 5 |
| 3 | 4 | 34 |
| 4 | 1483 | 1645 |
| 5 | 29142 | 29904 |
| 6 | 36554 | 38188 |
| 7 | 23456 | 26030 |
| 8 | 37760 | 43168 |
| 9 | 266518 | 273884 |
| 10 | 86293 | 90330 |
| 11 | 70711 | 73831 |
| 12 | 11631 | 13634 |
| 13 | 13274 | 16547 |
| 14 | 33423 | 36563 |
| 15 | 14304 | 18313 |
| 16 | 28324 | 32969 |
| 17 | 36157 | 40067 |
| 18 | 25838 | 29471 |
| 19 | 21778 | 24803 |
| 20 | 12481 | 18316 |
| 21 | 35809 | 41815 |
| 22 | 55893 | 61346 |
| 23 | 73479 | 78639 |
| 24 | 22436 | 25894 |
| 25 | 12376 | 18270 |
| 26 | 26294 | 33074 |
| 27 | 26196 | 30911 |
| 28 | 18975 | 22412 |
| 29 | 14126 | 17257 |
| 30 | 14296 | 20316 |
| 31 | 27865 | 33444 |
| 32 | 19812 | 23427 |
| 33 | 9882 | 12138 |
| 34 | 7820 | 8502 |
| 35 | 2370 | 2715 |
| 36 | 2311 | 2444 |
| 37 | 545 | 577 |
| 38 | 94 | 106 |
| 39 | 65 | 65 |
| Total: | 1,119,775 | 1,241,055 |

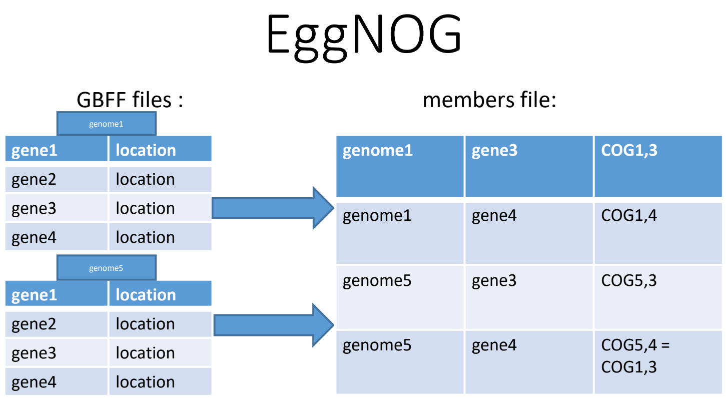


| Level | #Leaves | #Nodes |
|---|---|---|
| 3 | 0 | 1 |
| 4 | 2 | 26 |
| 5 | 15 | 58 |
| 6 | 32 | 125 |
| 7 | 72 | 262 |
| 8 | 505 | 917 |
| 9 | 1976 | 2288 |
| 10 | 1027 | 1174 |
| 11 | 725 | 759 |
| 12 | 83 | 87 |
| 13 | 8 | 8 |
| Total: | 4445 | 5705 |

(1) (2) (3)

Figure 1: **NCBI and EggNOG data**: (**1**) The NCBI Taxonomy hierarchy levels (**2**) The EggNOG DB file structure (**3**) The EggNOG-induced NCBI hierarchy levels

The maximal number of children for a node in this subtree is 135, for the node named '1883'. After the pruning process, every node from the original 4445 creator group of nodes of the subtree, is a leaf, and these are the only leaves of the subtree.

#### 2.4 Calculating the Synteny Index Values

In order to create SI values as described in the main text, we processed all pairs of genomes. For each pair of taxa from this group of 4,445 taxa, we calculated an SI value. As opposed to the theoretical model, in which every gene us unique in a genome, in real life, several genes in a genome may belong to the same COG, hence the SI calculation procedure was extended as follows: the SI between two genomes becomes the average over pairs of genes sharing the same COG (i.e. orthologs), i.e. if genomes $A$ and $B$ contain $n_C(A)$ and $n_C(B)$ copies (genes) of COG $C$, SI for the copies of $C$ will be calculated $n_C(A) \cdot n_C(B)$ times. The distribution of the number of shared COGs between the $9,876,790$ pairs of genomes is depicted in Figure 2(1).

The above yielded the squared $4445 \times 4445$ *SI table* that can be found at suppMaterial/SITable.meg file. The distribution of the $9,876,790$ values of SI, is given in Figure 2(2) and its histogram in Figure 2(3).

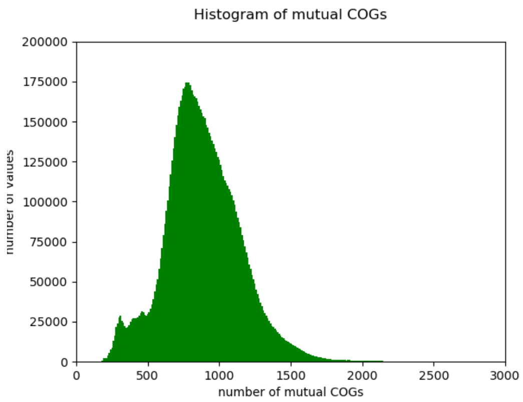


(1)

| Range | Num of Vals | Range | Num of Vals |
|---|---|---|---|
| 0.00-0.01 | 1230913 | 0.0-0.1 | 9800739 |
| 0.01-0.02 | 5864479 | 0.1-0.2 | 62364 |
| 0.02-0.03 | 1637858 | 0.2-0.3 | 10110 |
| 0.03-0.04 | 506036 | 0.3-0.4 | 2490 |
| 0.04-0.05 | 259418 | 0.4-0.5 | 880 |
| 0.05-0.06 | 139427 | 0.5-0.6 | 151 |
| 0.06-0.07 | 74519 | 0.6-0.7 | 12 |
| 0.07-0.08 | 43524 | 0.7-0.8 | 23 |
| 0.08-0.09 | 27638 | 0.8-0.9 | 21 |
| 0.09-0.10 | 16927 | 0.9-1.0 | 0 |

(2)

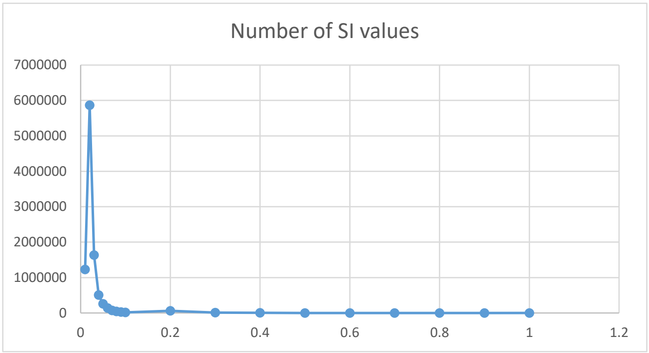


(3)

Figure 2: **Shared COGs and SI Values**: (**1**) Distribution of number of shared COGs between pairs of genomes (**2**) Distribution of SI values between pairs of genomes (**3**) Histogram of SI values between pairs of genomes

#### 2.5 Creating the SI trees:

The raw SI table served as input for three SI-based distance matrices, produced this raw table as follows:

1. **The** $1 - $ **SI**: Each entry $(i,j)$ in this table is obtained by $1 - [SI]_{i,j}$ For the first tree of the 3 - the distance between each taxa and each other taxa is 1-SI, when the SI value is the appropriate value for these two taxa. We named this tree '1-SI-tree'.

2. **The *heuristic-SI***: Here the distance between each pair of taxa is given by the heuristic formula, developed in [11] for $k = 10$ (see page 9, eq. 17 therein) where $n$ is the mean length of the two genomes whose mutual distance we calculate, and SI value taken from the raw SI matrix:
$$\hat{d} = \lambda t = -\frac{n}{3 - \frac{5k}{n-1}} \log(1 - \frac{1 - SI}{1 - \frac{2k}{n-1}}) \tag{7}$$

3. **The *exact-SI***: Here the distance represents the expected number of jumps as developed in the main text, which is the main contribution of the current work. The distance measure $d$ between each pair of taxa is obtained by developing the symbolic expression for $q_k$ for the specific value of $k = 10$ used for calculating SI. $d$ is subsequently obtained by solving numerically the equation:

$$\begin{aligned} exp(2*d)*(d+1)^{19}*SI = {} & 10*d^{18} + 90*d^{17} + 645*d^{16} + 2580*d^{15} + 8604*d^{14} + 20076*d^{13} \\ & + 39804*d^{12} + 59706*d^{11} + 76502*d^{10} + 76502*d^{9} + 64998*d^{8} + 43332*d^{7} \\ & + 24108*d^{6} + 10332*d^{5} + 3552*d^{4} + 888*d^{3} + 162*d^{2} + 18*d + 1 \end{aligned} \tag{8}$$

As tree reconstruction method, we chose *Neighbor Joining* (NJ) [9], a classical and robust method for reconstructing unrooted trees from pairwise distances. The distance to the reference NCBI taxonomy is measured using the classic *Robinson-Foulds* (RF) [8] symmetric difference distance. RF counts the number of internal edges in *exactly* one of the trees (and hence denoted symmetric difference) and consequently bounded by $2n - 6$ for two binary trees with $n$ leaves. The distances appear in the main text.

#### 2.6 Coloring Trees:

In order to produce intuitive sense of tree similarity we used the notion of color convexity [7, 6]. We partitioned the NCBI tree over the 4445 leaves into disjoint subtrees such that each subtree will have between 80 and 800 offsprings. Note that such a partition is not necessarily a cover, i.e. not all leaves are assigned to a subtree in the partition. This process yielded a partition of 14 subtrees over 4710 nodes (out of the 5705). We then associated each subtree with a unique “color”. This color is assigned to the leaves of that subtree (in the NCBI tree) and each leaf retains its color also under other trees. The (arbitrary) colors for these subtrees (RGB format), and the size of these subtrees are in table 1.

| Color | size |
|---|---|
| ('#ff0000') | 179 |
| ('#ff00ff') | 195 |
| ('#993300') | 198 |
| ('#ff9900') | 81 |
| ('#0033cc') | 123 |
| ('#009999') | 741 |
| ('#00ff00') | 313 |
| ('#66ffff') | 243 |
| ('#ff99cc') | 689 |
| ('#ffff99') | 480 |
| ('#6600ff') | 80 |
| ('#999966') | 532 |
| ('#990099') | 768 |
| ('#99cc00') | 88 |



Table 1: coloring subtrees

### References


[1] L. J. Allen. *An introduction to stochastic processes with applications to biology.* Chapman and Hall/CRC, 2010.

[2] W. J. Anderson. *Continuous-time Markov chains: An applications-oriented approach.* Springer Science & Business Media, 2012.

[3] W. Feller. *An introduction to probability theory and its applications*, volume 2. John Wiley & Sons, 2008.

[4] G. Grimmett, G. R. Grimmett, D. Stirzaker, et al. *Probability and random processes.* Oxford university press, 2001.

[5] S. Karlin and J. L. McGregor. The differential equations of birth-and-death processes, and the stieltjes moment problem. *Transactions of the American Mathematical Society*, 85(2):489–546, 1957.

[6] S. Moran and S. Snir. Efficient approximation of convex recolorings. *Journal of Computer and System Sciences (JCSS)*, 73:1078–1089, 2007. Earlier version appeared in APPROX/RANDOM 2005.

[7] S. Moran and S. Snir. Convex recolorings of strings and trees: Definitions, hardness results and algorithms. *J. Comput. Syst. Sci.*, 74(5):850–869, 2008.

[8] D. F. Robinson and L. R. Foulds. Comparison of phylogenetic trees. *Mathematical biosciences*, 53(1-2):131–147, 1981.

[9] N. Saitou and M. Nei. The neighbor-joining method: A new method for reconstructing phylogenetic trees. 4, 1987.

[10] C. L. Schoch, S. Ciufo, M. Domrachev, C. L. Hotton, S. Kannan, R. Khovanskaya, D. Leipe, R. Mcveigh, K. O'Neill, B. Robbertse, S. Sharma, V. Soussov, J. P. Sullivan, L. Sun, S. Turner, and I. Karsch-Mizrachi. NCBI Taxonomy: a comprehensive update on curation, resources and tools. *Database*, 2020, 08 2020. baaa062.

[11] G. Sevillya, D. Doerr, Y. Lerner, J. Stoye, M. Steel, and S. Snir. Horizontal Gene Transfer Phylogenetics: A Random Walk Approach. *Molecular Biology and Evolution*, 37(5):1470–1479, 12 2019.
